# Evidence or Confidence: What is really monitored during a decision?

**DOI:** 10.1101/2021.04.02.438275

**Authors:** Douglas G. Lee, Jean Daunizeau, Giovanni Pezzulo

## Abstract

Assessing our confidence in the choices we make is of paramount importance to making adaptive decisions, and it is thus no surprise that we excel in this ability. However, standard models of decision-making, such as the drift-diffusion model (DDM), treat confidence assessment as a post-hoc or parallel process that does not directly influence the choice, which depends only on accumulated evidence. Here, we pursue the alternative hypothesis that what is monitored during a decision is an evolving sense of confidence (that the to-be-selected option is the best) rather than raw evidence. Monitoring confidence has the appealing consequence that the decision threshold corresponds to a desired level of confidence for the choice, and that confidence improvements can be traded off against the resources required to secure them. We show that most previous findings on perceptual and value-based decisions traditionally interpreted from an *evidence-accumulation* perspective can be explained more parsimoniously from our novel *confidence-driven* perspective. Furthermore, we show that our novel *confidence-driven DDM* (cDDM) naturally generalizes to decisions involving any number of alternative options – which is notoriously extemporaneous using traditional DDM or related models. Finally, we discuss future empirical evidence that could be useful in adjudicating between these alternatives.

## Introduction

As humans, we are able to perform complex behaviors empowered by higher order cognitive processes, such as judgment and goal-directed decision-making. Beyond this ability, we are also able to monitor and assess such cognitive processes as they unfold. Known as metacognition, this feature of the human mind allows us to analyze the thoughts that we have and adjust them in a controlled manner so as to tune our own cognitive performance (Fleming et al., 2012). In decision-making, such metacognitive assessment comes in the form of choice confidence, or our subjective belief that what we chose was indeed the correct (or best available) option. Choice confidence is useful in that it can help us learn from our mistakes, or avoid a misallocation of scarce resources (Yeung & Summerfield, 2012). In decisions where there is an objectively correct answer (e.g., perceptual decisions), confidence is highly correlated with objective accuracy, although it is not always properly aligned (Fleming & Daw, 2017). It has been proposed that different individuals have different levels of metacognitive bias (a systematic mismatch between confidence and accuracy), metacognitive sensitivity (ability to discriminate between correct and incorrect responses based on feelings of confidence), and efficiency (sensitivity conditional on performance; (Fleming & Lau, 2014)). Moreover, in decisions where there is no objectively correct answer (e.g., preferential decisions), people are nevertheless able to report how confident they are about their choices (De Martino et al., 2013). At the neural level, various studies have mapped (retrospective and prospective) aspects of metacognitive ability to (lateral and medial) prefrontal areas and interoceptive cortices (see review in (Fleming & Dolan, 2012)). In spite of the extensive interest and progress in studies of confidence in different domains of decision-making (e.g., perceptual, preferential, economic), there is as of yet no widely accepted computational model that accounts for choice confidence. This is especially true in the field of preferential (value-based) choice, which is our primary interest. We believe that an acceptable contender can be generated from the class of models known as sequential sampling or accumulation-to-bound, and we introduce our contender model in this work.

Within the field of decision-making, accumulation-to-bound models are by far the most common^1^. Diffusion decision models (most notably, the drift-diffusion model or DDM) account very well for empirical data across a wide variety of domains, including perceptual and value-based decision-making (Forstmann et al., 2016). In value-based decision-making, the DDM simultaneously explains choice accuracy and response time (RT) distributions dependent on the relative values of the options that comprise a choice set (see review in (Ratcliff et al., 2016)). The basic premise of these models is that upon presentation of the choice options, information processing in the decision network of the brain provides a signal about which option is more valuable. This signal is assumed to represent some “true” value of the options, but because there is noise inherent in neural information processing, the signal is repeatedly probed until the system can reliably declare one option to be more valuable than the other. Figure 1 provides a simple illustration of the process for an example decision. The core of the DDM is an abstract decision variable that represents decision evidence (that one option is better than the other) as it accumulates across time. Though the evidence that this variable represents remains abstract in nature, there have been a variety of proposals as to what the evidence actually is: a function of the likelihood of each alternative response being correct, given the sampled information (Edwards, 1965; Laming, 1968; Stone, 1960); a comparison between sampled information and a mental standard (Link & Heath, 1975); a measure of strength of match between a memory probe and memory traces stored in long-term memory (Ratcliff, 1978); or the difference in spike rate between pools of neurons representing the alternative options (Gold & Shadlen, 2001).

**Figure 1:**
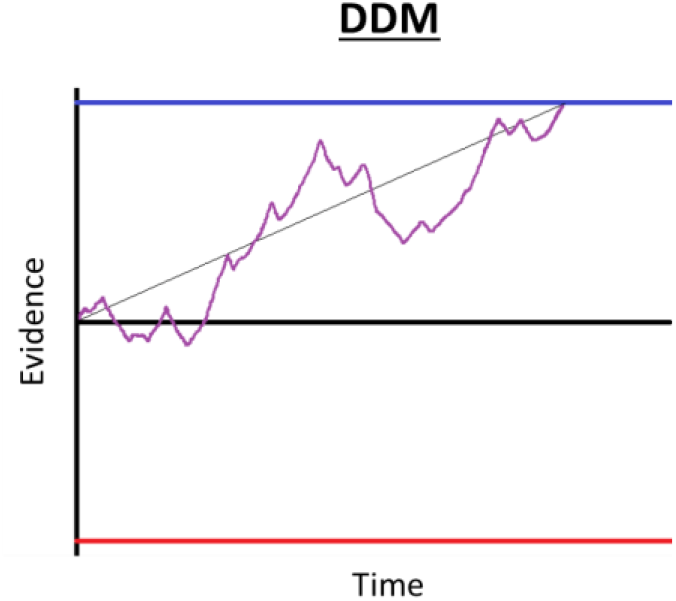
Basic drift-diffusion process illustration. Evidence (vertical axis) for one option versus the other (in a binary choice) evolves across time (horizontal axis) until a threshold is reached. The thin black line represents the ballistic trajectory, defined by the value of option 1 minus the value of option 2, scaled by the drift rate parameter. The purple trace represents the moment-by-moment accumulated evidence, which deviates from the ballistic trajectory according to random noise with variance set by the diffusion parameter. The blue and red lines represent sufficient evidence thresholds for options 1 and 2, respectively.

The basic components of the DDM, which relate to the accumulated evidence decision variable, are the *drift* rate or the trajectory at which it changes on average (proportional to the “true” value difference of the options), the *diffusion* noise parameter by which its trajectory is momentarily perturbed (the extent of signal corruption in the system), and the evidence *threshold* the arrival at which triggers a choice (the minimum level of evidence required to declare that one option is better than the other). It has been proven that the DDM can provide the optimal solution to simple decision-making problems, where prior beliefs about the option values are updated in a Bayesian manner through sequential evidence sampling until a choice can confidently be made based on posterior value beliefs (Fudenberg et al., 2018; Tajima et al., 2016). That line of work also shows that the optimal DDM decision thresholds should diminish over deliberation time^2^, in any situation where the difficulty of the task is not known *a priori* and where there is some cost associated with evidence accumulation (as would be the case in most preferential choice contexts; but see (Malhotra et al., 2018) for other scenarios where diminishing bounds might not be optimal). As the accumulator is here defined as the *difference* in the posterior estimates of option values, the height of the threshold at the time of choice could directly provide a measure of *choice confidence* (i.e., the probability that the chosen option was better (Pouget et al., 2016)). In this case, the collapse of the threshold as time lingers could be interpreted as a tradeoff between accepting a lower-than-desired level of evidence (and eventually, confidence) about the choice in exchange for a conservation of deliberation resources (e.g., time, metabolites, neural capacity).

One problem with using the threshold height as a direct readout of choice confidence is that it does not take into account the *precision* of the posterior value estimates. In the basic DDM, value estimates are point estimates, and the accumulating evidence does not consider the uncertainties of these estimates. A standard measure of uncertainty for (Gaussian) probability distributions is the variance of the distribution, or its inverse precision. The issue here is that with flat priors and Bayesian sequential updating, the difference in the posterior means of one pair of options could be equivalent to that of another pair, while each pair could have a different level of precision. For example, the posterior estimates of one option pair could have been formed from relatively inconsistent evidence samples (i.e., many samples in support of either option, hence lower-precision posteriors), whereas the posterior estimates of the other option pair could have been formed from relatively consistent evidence samples (i.e., most samples in support of the same option, hence higher-precision posteriors; see Figure 2). In this case, both choices might reach the same evidence threshold, but it seems intuitive that the option pair with the higher posterior precision should correspond to higher choice confidence. In fact, under the DDM framework, higher variability in the evidence accumulation process causes the process to terminate sooner (on average; see (Lee & Usher, 2021)), thus implying that less precise information would lead to higher confidence (because of the collapsing bounds). Using a threshold based solely on the value difference of the options as a measure of confidence cannot account for a fundamental psychological dimension of confidence – people are more confident when they decide based on more precise information (Lee & Coricelli, 2020; Lee & Daunizeau, 2020, 2021). To rectify this, the threshold height at the moment of the choice response would need to be transformed as some function of the precision of the value estimates (e.g., scaled by precision and passed through a sigmoidal function) in order to calculate confidence. However, although this might solve the problem mathematically, it would remain unclear as to why the deliberation process would seek a target (threshold) that would sometimes end up registering as high confidence and sometimes low (setting aside the issue of the eventual threshold collapse). More specifically, such an apparatus would allow the process to terminate even if the value estimates had very low precision, which would yield a low *post hoc* confidence readout (suggesting that deliberation had been terminated prematurely). It is unclear why the decision apparatus would sometimes make such low confidence choices by design. It seems more reasonable to believe that the same (initially high) level of confidence would always be the default target for similar decisions, and that the threshold that terminates the deliberation process should directly take into account the precision of the value estimates.

**Figure 2:**
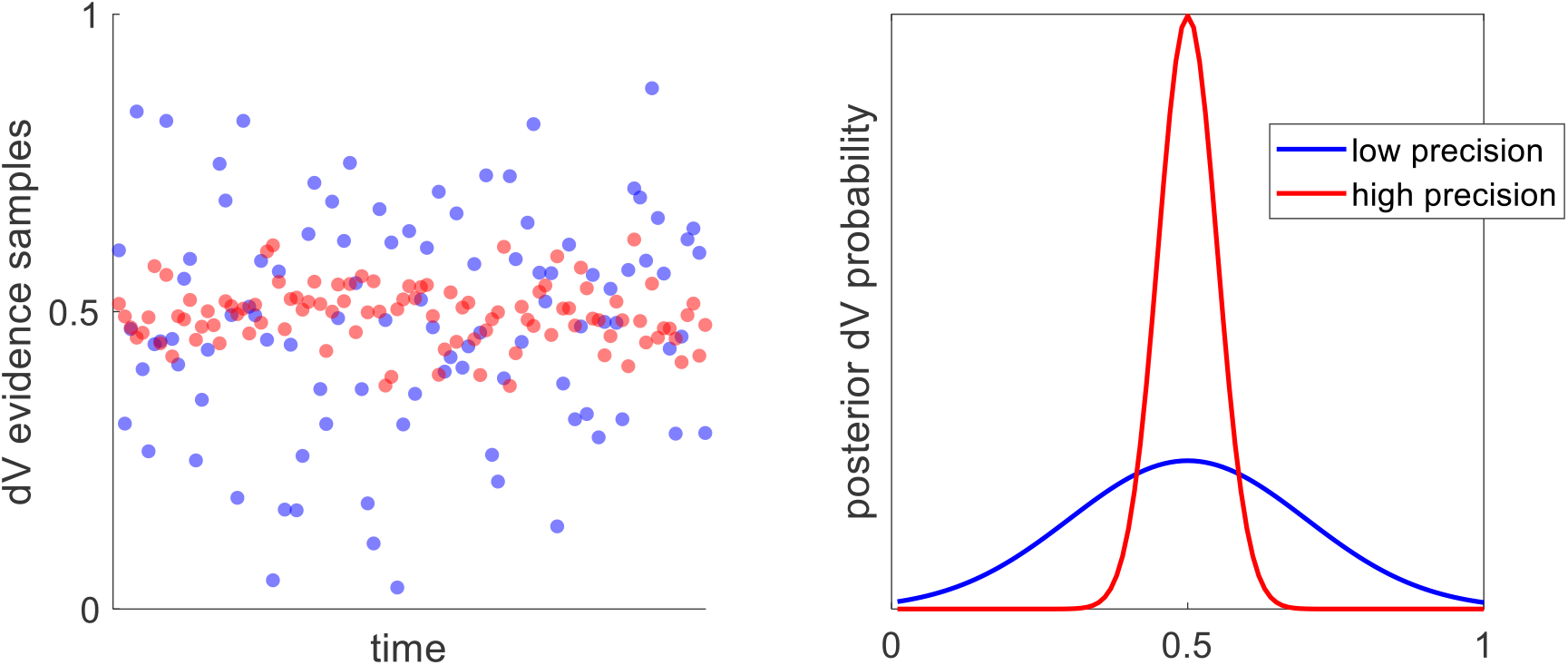
An illustration of how value estimates could have the same mean but different precision. Posterior estimates of the difference in option values (dV) might have similar means but very different levels of precision. The left panel shows sample streams of evidence for two different choice trials (one in blue, one in red). Notice that the samples fluctuate around the same average value, but with more (blue) or less (red) variability. The right panel shows what the posterior dV estimates for the two trials would look like—both have the same mean, but the red estimate has a much higher precision than the blue estimate. This higher-precision estimate should instill greater choice confidence.

Recent work has attempted to use the DDM to predict choice confidence in addition to choice probability and RT (Calder-Travis et al., 2020; Drugowitsch et al., 2019; Kvam & Pleskac, 2016; Moran et al., 2015; Moreno-Bote, 2010; Pleskac & Busemeyer, 2010; Yeung & Summerfield, 2012). However, most such attempts have required additional assumptions and components on top of the DDM itself, which detracts from the elegance of the original model and its ability to so reliably account for key decision variables. Given the success of the DDM in accounting for other key aspects of choice (i.e., choice probability and RT), and assuming that confidence about a choice is inherently linked to those other aspects (Kiani & Shadlen, 2009; Van Den Berg et al., 2016), one might conclude that the DDM should be able to simultaneously account for all three variables without the need for any *ad hoc* features. Calder-Travis et al (Calder-Travis et al., 2016) conducted a thorough analysis of various sorts of DDM found in the literature, specifically focused on their ability to predict choice confidence in a variety of datasets. However, those authors chose not to perform their model comparison while simultaneously accounting for choice, RT, and confidence. Furthermore, they only examined data from perceptual decision studies, so it is unclear if their findings generalize to preferential decisions.

One of the most frequently cited evidence accumulation models of confidence is the *two-stage dynamic signal detection* model (Pleskac & Busemeyer, 2010). As the name implies, this model includes two separate (but linked) stages: one in which evidence supporting the choice of one option versus the other is accumulated until a threshold is reached and the choice is made; one in which additional evidence that accumulates after the choice is made informs an estimate of confidence about that choice. Thus, confidence under this model is calculated only after the choice has been finalized, and it critically depends on the assumption that additional evidence continues to accumulate (via the same accumulator) after the choice response. Van den Berg and colleagues used a similar approach, and additionally suggested that confidence could be solicited at different time points (Van Den Berg et al., 2016). Those authors relied on the assumption that the evidence accumulator variable could be queried both at the time that the choice was reported as well as after some further delay to explain apparent changes of mind (i.e., participants later opted to choose the option that they initially rejected) and changes of confidence (i.e., participants later opted to choose the same option that they initially chose, but with a different level of confidence). This study therefore exposed the idea that not only does choice evidence accumulate across time, but confidence does as well – indeed, they are assumed to be based on the exact same accumulator variable. The idea that confidence can be monitored across deliberation time was further developed by Drugowitsch and colleagues, who suggest that the optimal stopping rule (i.e., the location and form of the response threshold) was determined by a comparison of the cost of continued evidence accumulation and the gain in confidence that the decision-maker should expect from continued evidence accumulation (Drugowitsch et al., 2012). While previous studies such as these already suggested that confidence was derived from the same information as the choice itself, that it could be queried at any point in time, and that such queries should optimally control the decision process (or at least its termination), they nevertheless all rely on the assumption that both the accumulator variable and the corresponding response thresholds pertain to raw evidence about the choice options. Assessments of confidence, accordingly, would require that the evidence signal be further processed or transformed.

Here, we pursue an alternative hypothesis that the response threshold in the DDM should directly represent a target level of choice confidence, and thus that the evolving decision variable should represent the momentary level of confidence across time. This idea has qualitatively been proposed before as the diminishing criterion model for metacognitive regulation of time investment (Ackerman, 2014). In the present work, the decision variable is quantitatively defined as a sigmoidal transformation of the absolute difference in posterior value estimates scaled by the posterior precision (i.e., the normative posterior probability that one option is better than the other, given the accumulated evidence thus far). Note that this allows for the posterior precision to be incorporated dynamically throughout decision deliberation. The underlying computational architecture can be conceptually divided into two modules. One module monitors confidence changes and releases response inhibition when a satisficing target confidence level has been achieved (see (Lee & Daunizeau, 2021)). The other module enables the choice response in favor of the option deemed to hold the highest value. This seemingly superficial modification of the basic DDM has three main implications. First, whereas previous models have assumed that choice confidence is read out as a transformation of the accumulated evidence after the choice has been made, we propose that the transformation from evidence to confidence occurs during (and is indeed an integral component of) the decision process (Drugowitsch et al., 2012; Tajima et al., 2016). We note that this does not preclude post-decisional evidence from being incorporated into post-decisional confidence reports (Desender et al., 2021; Moran et al., 2015; Murphy et al., 2015; Navajas et al., 2016; Pleskac & Busemeyer, 2010). Second, the ensuing decision process gracefully generalizes beyond two-alternative decisions. This is because confidence (i.e., the subjective probability that the option estimated to be best at any point in time will actually yield the highest value) can be derived for any number of alternative options (see Appendix A). Third, the process thus signals when a choice can be made with a satisfactory level of confidence, but it does not directly control *which* option will be selected.

Our formulation addresses a number of known shortcomings of existing sequential sampling models. First, as we hinted at above, basic DDMs cannot account for the impact of value estimate uncertainty. Decision-makers form not only estimates of value for the different options that they consider, but also assessments of (un)certainty about those value estimates (De Martino et al., 2013; Gwinn & Krajbich, 2020; Lee & Coricelli, 2020; Lee & Daunizeau, 2020, 2021; Polanía et al., 2019). These feelings of certainty are important, because they alter choice behavior (Lee & Coricelli, 2020; Lee & Daunizeau, 2020, 2021). Most existing versions of the DDM (or any other sequential sampling model) exclude the possibility that option-specific value certainty might play a role in the decision process. However, it has recently been shown that the DDM indeed provides a better explanation of choice data when the drift rate is adjusted to reflect the different degree of certainty that the decision-maker has about the value of each option (Lee & Usher, 2021). The authors demonstrated that the so-called *signal-to-noise DDM* (snDDM) is capable of accounting for the positive impact of option-specific value certainty on choice consistency and confidence, and the negative impact on RT (Lee & Coricelli, 2020; Lee & Daunizeau, 2021). The snDDM presents an advance in sequential sampling models of choice, but it nevertheless relies on evidence samples drawn from static distributions. Our formulation, on the contrary, is driven by value estimates that evolve across deliberation. In particular, our formulation fundamentally includes an increase in value certainty during choice deliberation, which is consistent with empirical findings that value certainty ratings are generally higher after choices compared to before (Lee & Coricelli, 2020; Lee & Daunizeau, 2021).

### An illustrative example of the confidence-driven DDM (cDDM)

Our proposed *confidence-driven DDM* (cDDM) offers several advantages over the standard DDM. For one, the cDDM could be more easily adapted to decisions between any number of options; standard DDMs only apply to binary decisions (but see (Kvam, 2019)). Furthermore, the cDDM could be directly applied to potentially any type of decision without having to adjust the meaning of its components or substantially alter its parameters (e.g., perhaps the target confidence level for a decision-maker might be similar across decision domains, whereas the relative evidence requirements might grossly differ). Finally, from a computational perspective, it would seem more useful (or parsimonious) for the brain to assess the reliability of the relevant information (i.e., confidence about which option seems better) as a choice response freely develops, rather than simply keeping a tally of how many stochastic samples favor each option, stopping when some arbitrary threshold has been reached, and only then checking to see how confident it is about the choice that would have already been made by that point. Our proposal is in line with previous work showing that decision-makers often change their minds about which option is better during the course of deliberation, and that such changes of mind coincide with changes of confidence (Van Den Berg et al., 2016). Although that previous work only focused on changes of confidence that occurred between response onset and response termination (in that task, participants registered their responses by moving their hands from the center of the screen to one of the four corners), it directly implies that confidence is monitored across time as long as evidence continues to accumulate. This suggests that confidence is indeed continuously monitored throughout deliberation, with (usually unobservable) changes of mind being triggered whenever confidence (that the current to-be chosen option is the best) disappears.

In addition, the cDDM includes a “collapsing bound” mechanism, which encourages the decision process to terminate even if the target level of confidence has not yet been achieved – and hence to make the choice with a lower level of confidence, because further resource expenditure (cognitive, metabolic, or other) to increase confidence is deemed too costly to continue (see (Lee & Daunizeau, 2021)). This is analogous to what happens in the optimal DDM, where the threshold collapses explicitly due to a cost of deliberation that accumulates over time ((Fudenberg et al., 2018; Tajima et al., 2016); see (Malhotra et al., 2018) for an alternative rationale for collapsing thresholds). An alternative (but mathematically similar) perspective is that the decision mechanism includes an urgency signal that amplifies the incoming decision evidence, which (akin to the collapsing bound mechanism) encourages an end to deliberation and a final choice response without excessive further delay (Churchland et al., 2008; Thura & Cisek, 2014; van Maanen et al., 2016).

Thus, the decision process of the cDDM can be summarized as a continuous readout of choice confidence, driven by a continuously-updated value signal (both estimate and precision for each choice option) towards a predetermined target confidence level, which is gradually reduced in consideration of the growing resource expenditure or urgency of the decision (see Figure 3 for an illustrative example).

**Figure 3:**
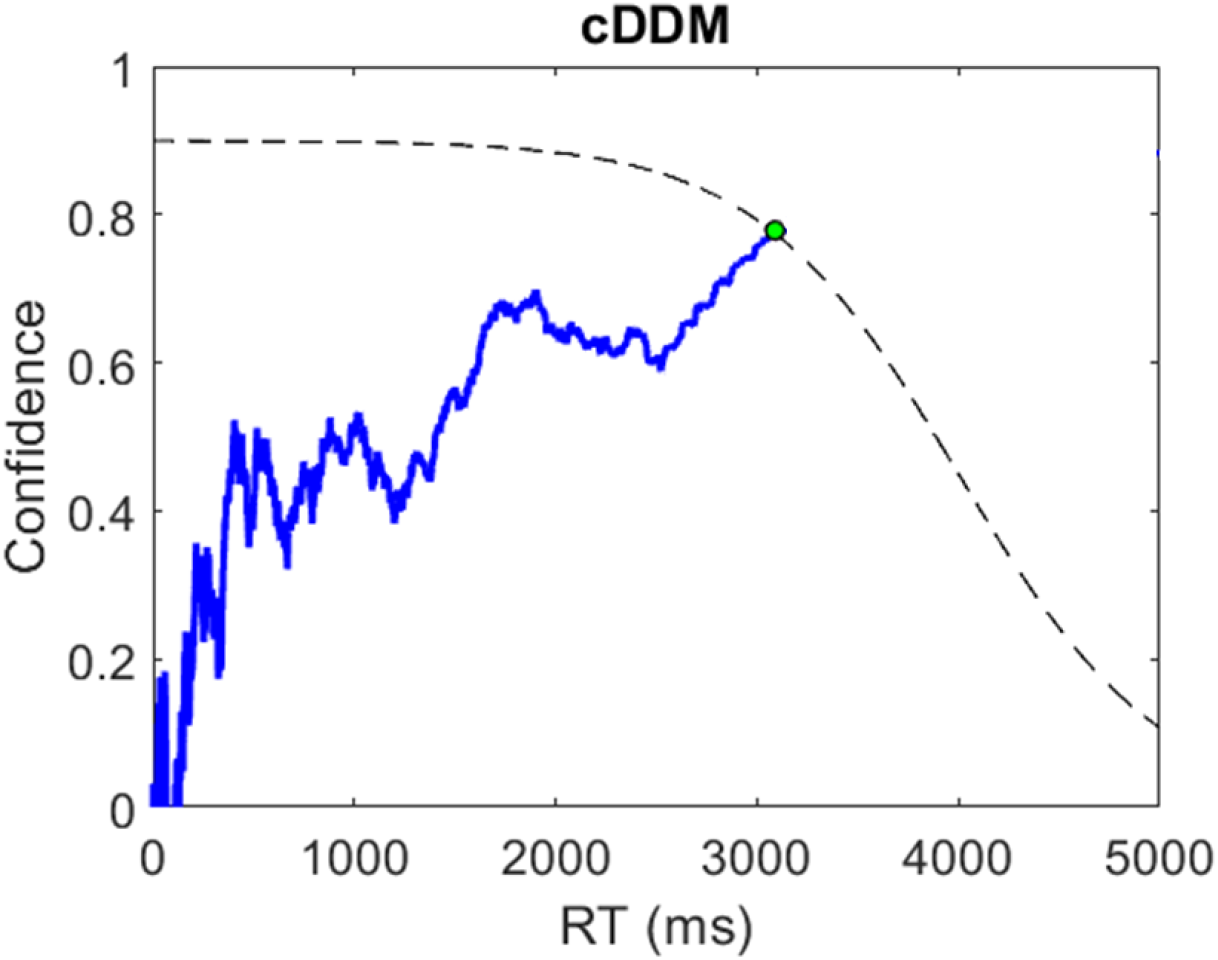
An illustrative example of the cDDM process. The blue trace illustrates the evolution of choice confidence across response time (RT). At t=0, the options are unknown, and thus confidence = 0. Momentary precision-weighted value evidence drives confidence until it reaches a threshold and a choice is made. Note that the target confidence threshold collapses over time, as the decision-maker recognizes the importance of conserving resources (e.g., time and cognitive effort) rather than continuing to increase confidence.

## MODEL

In this paper, we propose a novel version of the drift-diffusion model (DDM) that involves only minor alterations to the core mathematics, but major alterations to the core conceptual interpretation of decision control. Taken together, these alterations allow our new cDDM to directly account for choice confidence alongside choice consistency and response time (RT), in an integrated manner. Below, we first summarize the standard mathematical formulation of the basic DDM, complete with some well-known optional parameters (starting point bias, non-decision time, collapsing bounds). We then describe the alterations that distinguish the cDDM from the traditional DDM, as well as their contrasting assumptions and interpretations.

In the basic DDM, the decision deliberation process begins with both time and cumulative evidence equal to zero. At each time step (t) after the onset of deliberation, evidence (e) for each choice alternative (i) is drawn from an independent Gaussian distribution whose mean (μ_i_) is the true value of the option and whose variance (σ^2^) represents the amount of signal corruption due to the inherent stochasticity of neural processing (which is the same for all options):

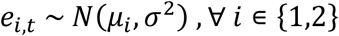

The evidence for each option at each time step is compared and the balance is added to the cumulative total evidence (x) for one option over the other:

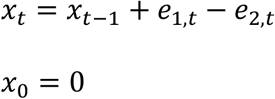

The deliberation process proceeds until the accumulated evidence reaches one of two symmetric predetermined threshold bounds (b for option 1, -b for option 2, b>0). It is possible to bias the starting point of the process (x_0_) away from zero, but this is arbitrary unless there is a good reason to assume that one option is *a priori* more likely to be better (e.g., instruction about or observation of an asymmetrical environment during a sequential decision-making task). It is also possible to include a measure of non-decision time (NDT) to represent the strictly positive time that it takes for the brain to process perceptual information while recognizing the options or motor control information while entering a response, which is simply added to deliberation time (DT) to calculate total RT.^3^ As mentioned above, one possible way to map from evidence to confidence in the traditional DDM would be to pass it through a sigmoidal transformation. Figure 4 illustrates simulated evidence accumulation, and the corresponding confidence transformation, for 100 trials of varying difficulty (i.e., unsigned value difference between the options).

**Figure 4:**
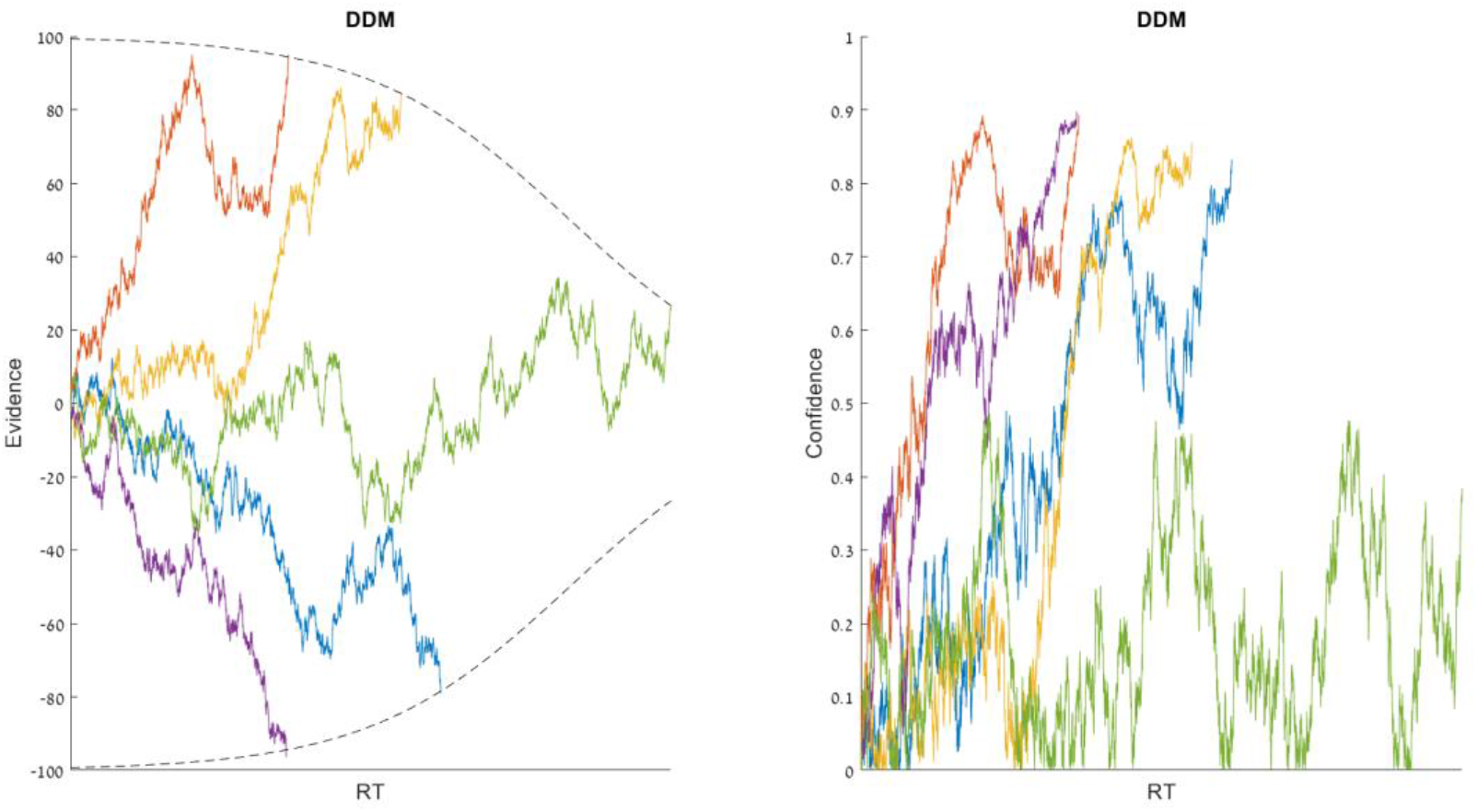
Simulated DDM trials. An illustrative example of five simulated trials of varying value difference (dV), based on a basic DDM with collapsing bounds. Each colored curve represents one trial. The left plot shows the accumulation of evidence across time, where each process terminates upon reaching one of the bounds. The right plot shows how the evolution of confidence would look if it were a moment-by-moment sigmoidal transformation of the cumulative evidence.

In the cDDM, the momentary variable that is monitored is not merely a relative value signal for the options (as in the basic DDM), but rather an assessment of confidence (c_t_) that considers both the momentary value estimates of each option (v_i,t_) and the momentary precision of those value estimates (p_i,t_), passed through a sigmoidal transformation to compute the probability that one option is better than the other:

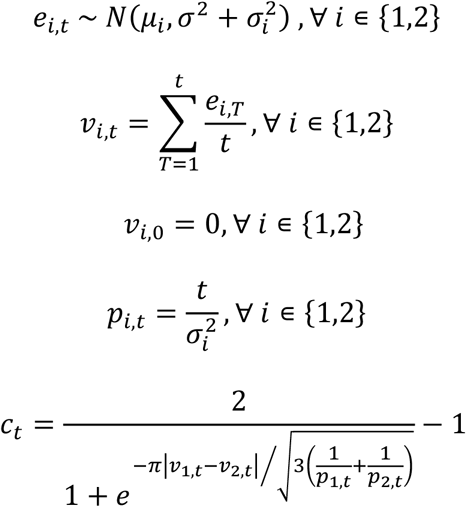

where now each option has its own specific inverse precision term (σ^2^_i_) in addition to the generic processing noise (σ^2^), and the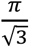term in the confidence equation derives from a moment-matching approximation to the Gaussian cumulative density function (Daunizeau, 2017a). Anecdotally here, the standard sigmoid (logistic) function undergoes an affine transformation to ensure that the confidence readout falls within the range [0,1), with 1 indicating maximal confidence about knowing which option is correct and 0 indicating a complete lack of confidence. Note that the evidence (e) that lies at the core of the process is the same as in the standard DDM, namely momentary signals about the relative values of the choice options (e.g., neural firing rates in pools of neurons representing either option). This enables a direct comparison between the standard DDM and the cDDM, in terms of numerical simulations. Note that the update of the sufficient statistics of value representations can be described in two distinct manners, including as a Markov decision process (which involves simple difference equations).

The decision process will continue to step forward in time and acquire additional samples of value evidence until confidence reaches the boundary (b) representing a predetermined target confidence level. The final inferred values of the options will be v_i_ ^τ,^ where τ is the time at which the process terminated. The choice will be implemented in favor of whichever option corresponds to the highest value of v_i_ ^τ,^ (i.e., response inhibition will be released when the confidence threshold is crossed, allowing the strongest value signal at that point to drive the decision-maker’s response behavior). Figure 5 illustrates simulated confidence accumulation, and the corresponding value estimate refinement, for 100 trials of varying difficulty.

**Figure 5:**
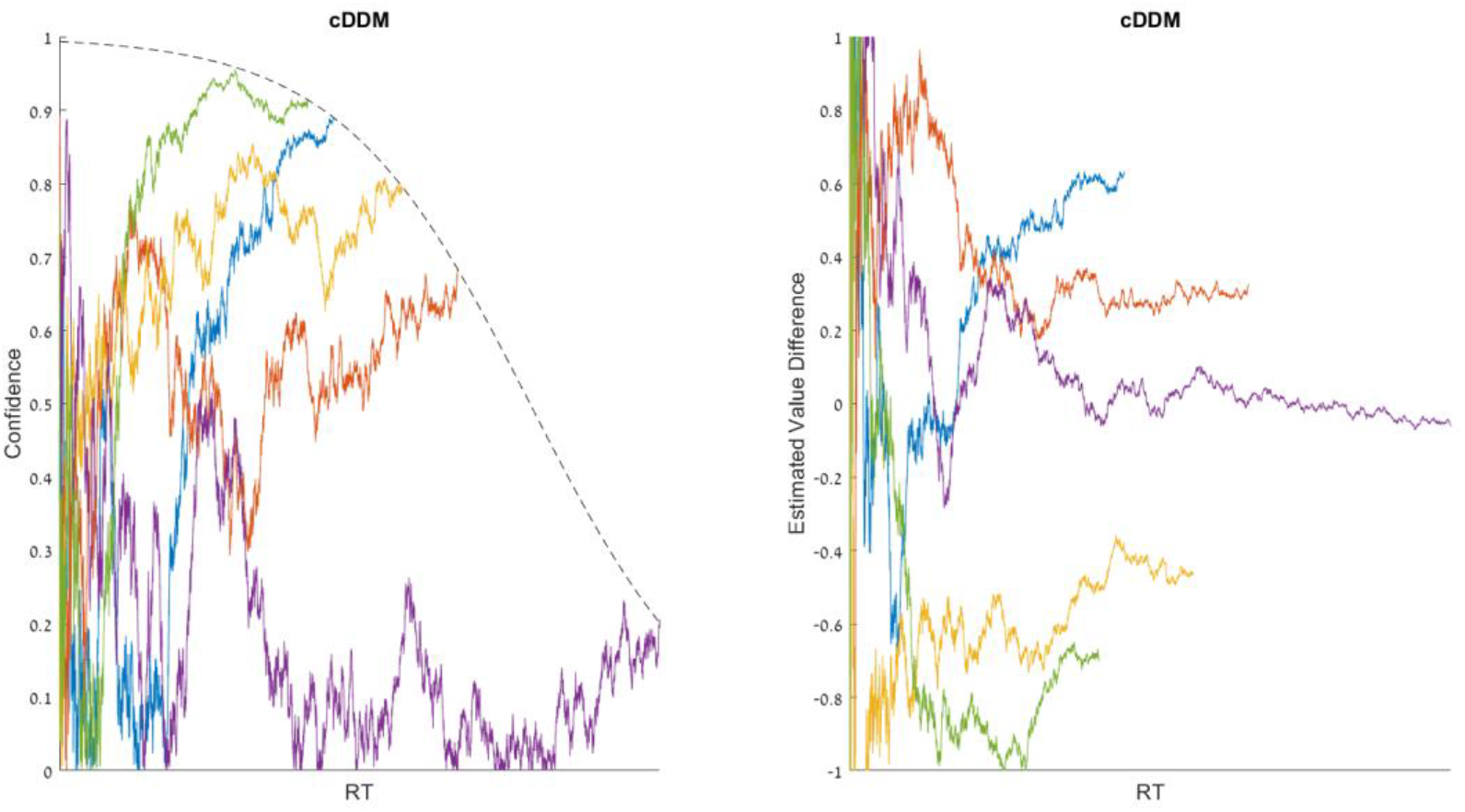
Simulated cDDM trials. An illustrative example of five simulated trials of varying value difference (dV), based on the cDDM. Each colored curve represents one trial. The left plot shows the accumulation of confidence across time, where each process terminates upon reaching the bound. The right plot shows how the evolution of estimated value difference would look if it were to be extracted from the confidence signal.

## RESULTS

Although we present the cDDM here as a theoretical model, it is nevertheless important for us to demonstrate that it is able to account for certain relevant findings previously reported in the literature. To permit this, we simulated data under the cDDM. Specifically, we created 10,000 trials each with two options whose means were independently drawn from a uniform (0,1) distribution and whose standard deviations were independently drawn from a uniform (1,2) distribution. To prevent outlier trials with excessively long response time (RT), we arbitrarily set a maximum RT at 10,000 time steps (t) at which point the accumulation process would stop regardless of the current state of confidence. We arbitrarily set the shape of the response threshold (t = 0:10,000) according to the formula: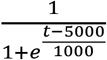. This established an initial (t = 0) target confidence of approximately 1, which gradually decayed with a rate that is at first increasing, then decreasing in approach of a target confidence of approximately 0 at t = 10,000. For the free parameters for the rate of evidence accumulation and the degree of neural processing noise, we manually selected parameters a value of 0.1 and 6, respectively, which seemed to produce output (choice probability, RT, and confidence) that reasonably resembled previously-published empirical data in terms of the observed range of each variable and the qualitative relationships between the variables. Varying the choice of parameters beyond a certain range could alter the qualitative nature of the simulated output. However, as the purpose of this theoretical study is merely to demonstrate the workings of the cDDM and its ability to qualitatively account for empirical findings, not to formally fit or quantitatively compare models, a proper formal parameter sensitivity analysis is beyond the scope of this work. As we present below, the cDDM can qualitatively reproduce a wide range of findings using a consistent set of parameters.

To simulate the cDDM process, each trial started with time, value estimates and precision for each option, and confidence all reset to zero. At each time step, a random sample of evidence was drawn for each option, which was scaled by the drift rate. The value estimate of each option was updated in a Bayesian manner, with the new evidence and already incorporated evidence scaled by their relative precisions before being summed together. The value precision for each option was updated as the previous level of precision plus the precision of the new evidence. Momentary confidence was calculated based on the momentary value estimates and precision, and the process terminated when confidence reached the threshold.

The most common behavioral findings, reported in a number of previous studies, are the general relationships between choice difficulty (inversely quantified as the difference in the choice option values as subjectively rated in a separate task, or dV), response time (RT), and choice confidence. Figure 6 presents these patterns as exemplified in data pooled together from three different studies (Lee & Coricelli, 2020; Lee & Daunizeau, 2020, 2021), alongside the patterns present in data simulated under the cDDM (where dV indicates the true latent value difference of the options on each trial). The cross-participant mean correlations between dV and RT, between dV and confidence, and between RT and confidence in the empirical data are -0.22 (*p* < .001), 0.31 (*p* < .001), and -0.30 (*p* < .001), respectively. In the exemplary set of cDDM parameters, the correlations are -0.75, 0.72, and -0.98. Note that without even needing to perform quantitative model fitting, it is evident that the cDDM qualitatively reproduces the same patterns found in the empirical data. The weaker correlations in the empirical data relative to the simulated data are expected due to additional sources of noise that are outside the scope of the model. Furthermore, the cDDM cannot reproduce the stochasticity in the empirical relationship between RT and confidence, because that relationship is explicitly defined in the model formulation. The origin of these relationships under the cDDM can be summarized as follows. Trials with lower dV will take longer to muster confidence, through either precision gain or estimate revision (thus, the negative relationship between dV and RT). These trials will reach the collapsing confidence threshold at a later point in time on average (thus, the positive relationship between dV and confidence). The negative relationship between RT and confidence is by construct, because the confidence threshold collapses across deliberation time.

**Figure 6:**
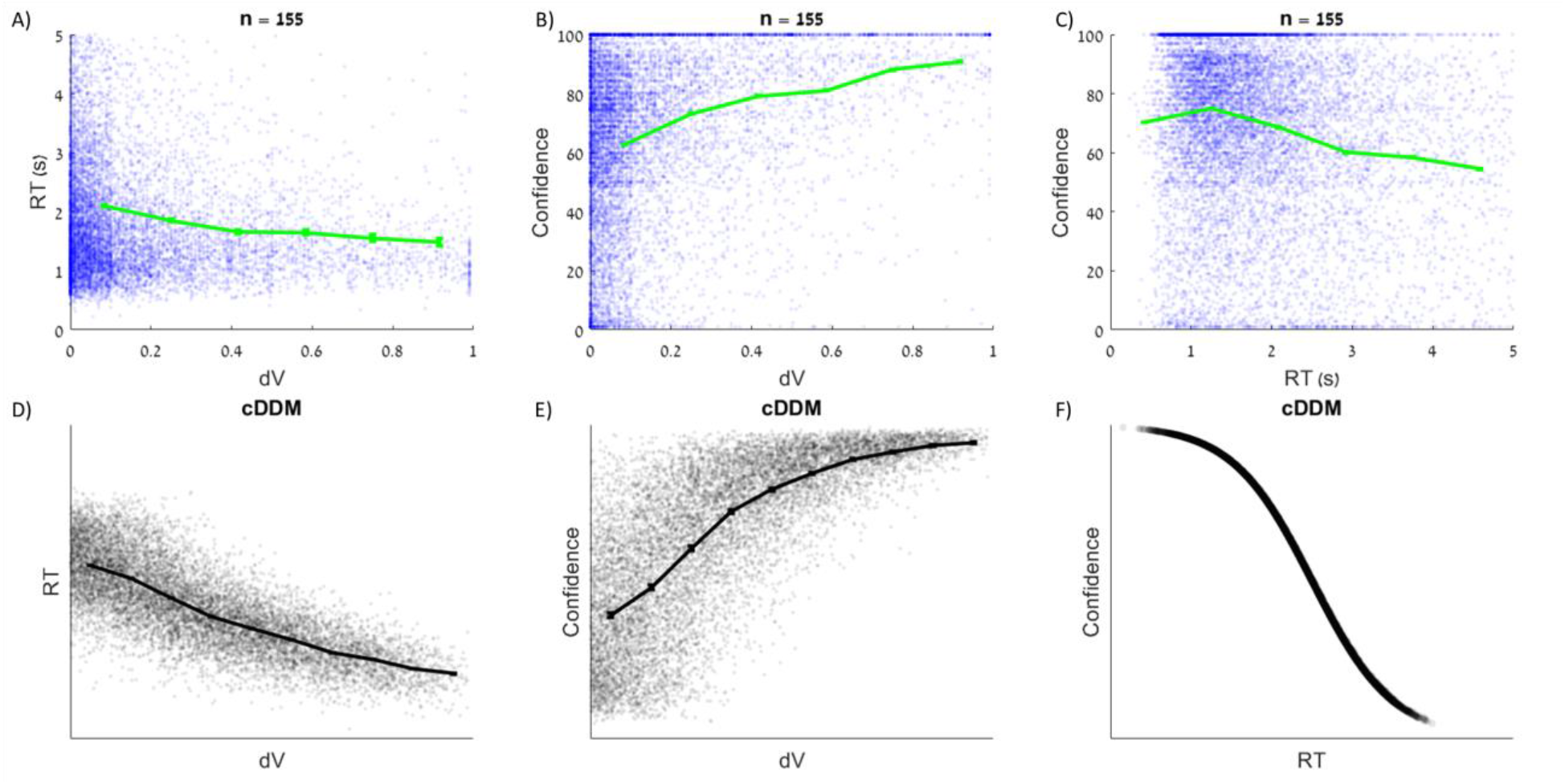
Relationships between choice ease, response time, and choice confidence. In experimental data pooled together from three previous studies (n=155), response time (RT) decreases with choice ease (unsigned value difference or dV; panel A), confidence increases with dV (panel B), and RT and confidence are negatively correlated (panel C). Each blue dot represents one trial; green curves show the means across equally-spaced dV bins. The same patterns were present in data simulated under the cDDM (panels D-F).

The three-way relationship between choice probability, response time, and choice confidence was summarized in a convenient format by de Martino and colleagues (De Martino et al., 2013). Specifically, the authors showed that choices were more consistent with subjective value ratings when participants were more confident about those choices, resulting in higher logistic regression slope parameters for more confident choices. They also showed that more difficult choices (i.e., lower dV) corresponded to longer deliberation (i.e., higher RT), and that more confident choices were made more quickly, even when accounting for difficulty level. We replicated those findings using the pooled data from the studies mentioned above (Lee & Coricelli, 2020; Lee & Daunizeau, 2020, 2021). In this empirical data, the cross-participant mean logistic slopes for low and high confidence trials were 6.6 and 15.0, respectively (difference in slopes = 8.4, *p* < .001; Figure 7A-B). Cross-participant mean RT for low dV / low confidence trials was greater than for high dV / low confidence trials (0.27s, *p* < .001), as well as for low dV / high confidence trials (0.49s, *p* < .001). Cross-participant mean RT for high dV / high confidence trials was lesser than for low dV / high confidence trials (0.35s, *p* < .001), as well as for high dV / low confidence trials (0.57s, *p* < .001). We also replicated the same findings in data simulated under the cDDM (Figure 7). The observation that choices are more consistent with value ratings on trials that had higher confidence can simply be explained by the fact that most high-confidence trials are easy (i.e., high value difference) and most low-confidence trials are difficult (i.e., low value difference). The three-way relationship between dV, confidence, and RT is more interesting. Under the cDDM, even when controlling for dV, trials will randomly vary in terms of how much congruent versus incongruent information is processed (i.e., how the trial-specific random information samples help or hinder momentary confidence). Some trials will benefit from more information earlier during deliberation, and thus terminate with higher confidence and lower RT. Other trials will not receive the same benefit, and thus terminate with lower confidence and higher RT.

**Figure 7:**
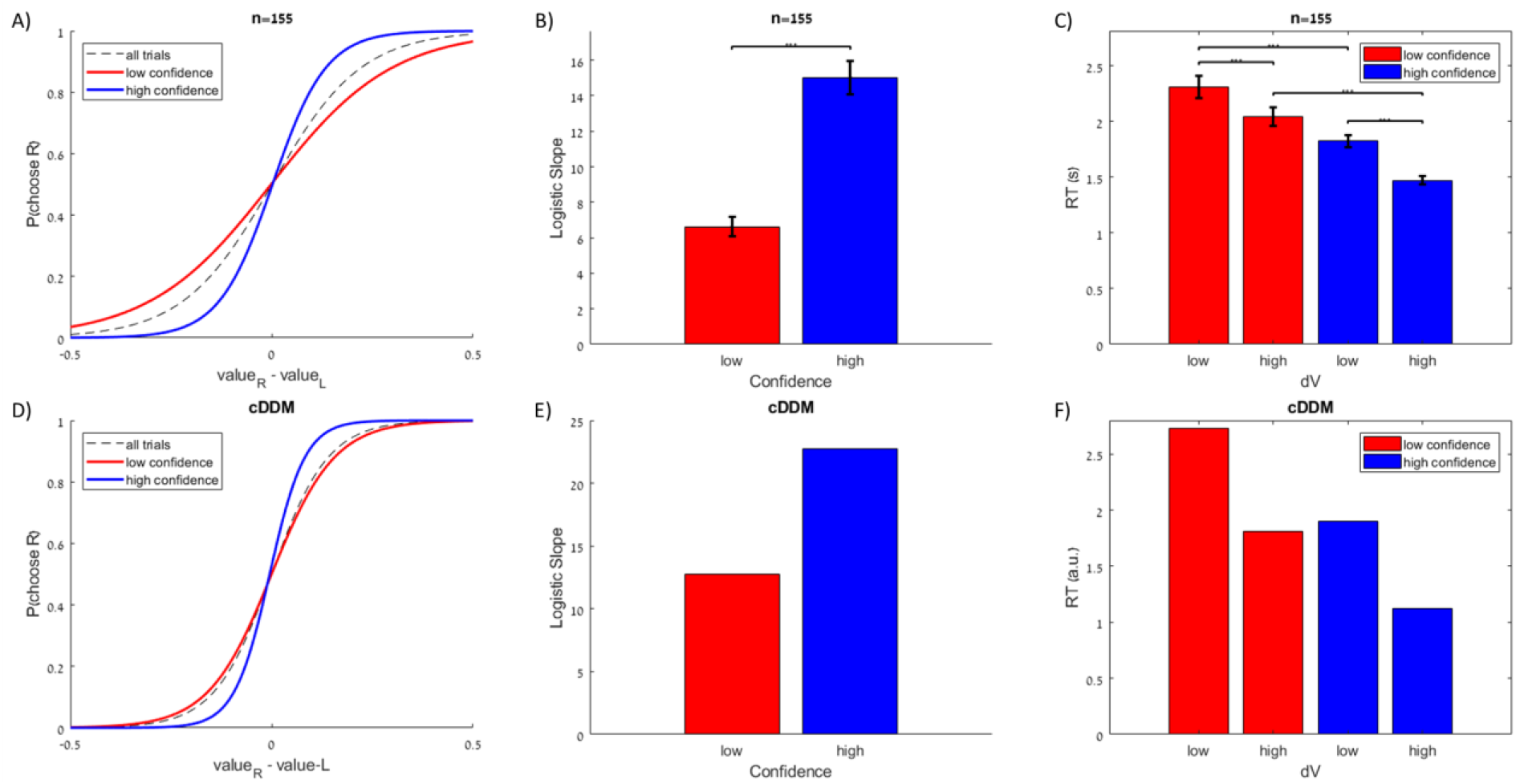
Confidence corresponds to greater choice consistency and lower response time. In experimental data pooled together from three previous studies (n=155), choices are more consistent with value ratings on high versus low confidence trials (panels A-B, blue versus red; median split on confidence, within participants). Responses are faster for high versus low dV trials (median split on dV, within participants), as well as for high versus low confidence trials within both high and low dV trials (median split on confidence within median split on dV, within participants; panel C). The same patterns were present in data simulated under the cDDM (panels D-F).

Another interesting finding reported in the literature is the observation that choice confidence increases with the value difference of the options (dV) for correct choices, but confidence decreases with dV for error choices. This was demonstrated in rats, which signaled their choice confidence by how long they were willing to wait in anticipation of their predicted reward in an olfactory discrimination task (Kepecs et al., 2008). It turns out that this pattern is also observable in the behavioral data from the studies mentioned above (Lee & Coricelli, 2020; Lee & Daunizeau, 2020, 2021), although it was not previously reported. Across participants, the mean correlation between dV and confidence for consistent and trials was 0.37 (*p* < .001), and the mean correlation between dV and confidence for inconsistent trials was -0.10 (*p* < .001; Figure 8). Data simulated under the cDDM also exhibit this pattern (correlation for consistent trials = 0.71, correlation for inconsistent trials = -0.09; Figure 8). Importantly, the same exact simulated data (in particular, the same generating parameters) were used to create all of the plots in Figures 6, 7, and This effect arises under the cDDM because inaccurate responses are caused entirely by noise that pulls the accumulator away from its pure (i.e., noise-free) ballistic trajectory. Such noise delays the choice response (in contrast to noise that pushes the accumulator towards an accurate response, which hastens the choice response). Therefore, the average response time for inaccurate trials will be longer than for accurate trials. Because the cDDM includes a confidence threshold that collapses over time, inaccurate trials will (on average) conclude with lower levels of confidence than accurate trials.

**Figure 8:**
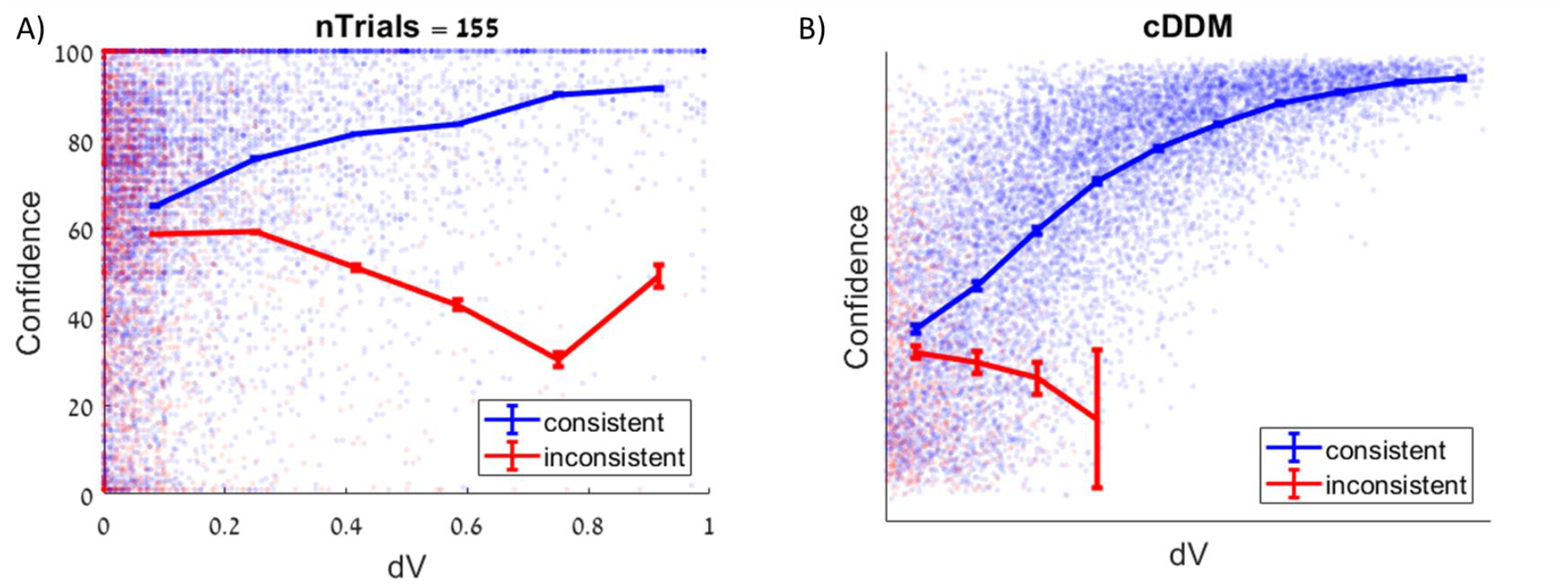
The relationship between choice ease and choice confidence is different for consistent and inconsistent trials. In experimental data pooled together from three previous studies (n=155), confidence increases with choice ease (unsigned value difference or dV) for consistent trials (where the chosen option aligns with value ratings; panel A in blue) but decreases with dV for inconsistent trials (panel A in red). Each dot represents one trial; curves show the means across equally-spaced dV bins. The same patterns were present in data simulated under the cDDM (panel B).

Finally, we wish to point out that in addition to replicating a variety of patterns across trials, the cDDM can also account for patterns observed within trials. As an example, consider the neural data reported in (Kiani & Shadlen, 2009). Of particular interest is the ramping firing activity of the lateral intraparietal cortex (LIP) neurons in rhesus monkeys as they decided in which direction the majority of displayed dots was moving on each trial. The authors show in their Figure 4 that the trajectories of the recorded neural activity match the evolution of the decision variable in their model, with activity rising towards a plateau on trials with strong supporting evidence and activity evolving in the opposite manner on trials with strong opposing evidence. However, in order to get the patterns in the neural data to match those predicted by their model, the authors first had to de-trend the neural responses by subtracting the mean response at each moment (Kiani & Shadlen, 2009). While it is simple to perform such transformations mathematically, it might not be so innocuous for the brain to perform them via neural information processing. We believe it is noteworthy that such neural response patterns would match those predicted by the cDDM (see Figure 5 above) without the need for any normalization. This can be taken as evidence that decision-related neural activity (in LIP) might align more directly with the decision variable in the cDDM (i.e., confidence) than the decision variable in the standard DDM (i.e., evidence).

### Empirical hurdles for confidence models to explain

The literature on choice confidence has exposed a variety of different empirical findings. Many of these findings are so robust that it has been proposed that any worthy model of confidence should be able to account for them (Pleskac & Busemeyer, 2010). We here put the cDDM to the test and note that it predominantly passes these hurdles:

#### 1. Speed-accuracy trade-off

It has long been known that decision time and accuracy are positively correlated, in the sense that participants respond more accurately (on average) when they are allowed to or otherwise choose to take more time before reporting their choices. Within the accumulation-to-bound framework, this is typically enabled by allowing the response boundary to vary across conditions: contexts where fast decisions are required or encouraged impel participants to set lower response boundaries compared to those in contexts where accurate decisions are encouraged. The cDDM is a specific instance of the accumulation-to-bound framework and is thus able to account for this speed-accuracy trade-off effect. Note, however, that this hurdle describes comparisons across different choice contexts. Within a given context, the both the basic DDM and the cDDM will predict the opposite pattern (i.e., longer average deliberation time should correspond to lower average accuracy).

#### 2. Positive relationship between confidence and stimulus discriminability

It has also long been known that confidence is positively correlated with stimulus discriminability (e.g., the difference in subjective value ratings between options in a value-based decision). Under the cDDM, confidence is defined as a monotonically-increasing function of value difference. Thus, while sampling stochasticity will lead to trial-by-trial variations in confidence levels at the time of choice, there will always be a positive correlation (across trials) between value difference and confidence.

#### 3. Resolution of confidence

Choice accuracy and confidence are both increasing functions of stimulus discriminability, otherwise referred to as choice ease. It is therefore not surprising that accuracy and confidence are themselves positively correlated. However, it has been shown that the positive relationship between accuracy and confidence remains even after controlling for choice ease/difficulty. In essence, this means that for a fixed difficulty level, participants report greater confidence for choices that turned out to be accurate than for choices that turned out to be inaccurate. We verified this via simulation (see Figure 8 above).

#### 4. Negative relationship between confidence and decision time

Many previous studies have reported a strong relationship between decision time and confidence, where slower decisions are generally associated with lower confidence. This effect is fundamental to the cDDM, where the confidence threshold (i.e., the boundary representing the momentary target confidence level at which a response will be made) decreases across decision time.

#### 5. Positive relationship between confidence and decision time

It has previously been shown that participants report higher levels of confidence (on average) during experimental conditions where they are either allowed or encouraged to take more time to deliberate before entering their choice responses. Under the cDDM, momentary confidence levels (during deliberation, prior to response) generally increases with decision time, as considering more information samples leads to higher-precision value estimates and thus higher choice confidence. The cDDM assumes that decision-makers will continue to deliberate until their confidence reaches a threshold level. However, if one were to relax this assumption and force participants to respond early (as is done in so-called interrogation paradigms), it should be clear that average confidence levels will be greater when participants are allowed more time before responding. Alternatively, if one were to retain the threshold feature of the cDDM and merely encourage shorter or longer deliberation times (as is done in speed versus accuracy optional stopping paradigms), it would also be the case that the slower conditions (i.e., those where accuracy was encouraged) would generally lead to greater confidence as compared to the faster conditions (i.e., those where speed was encouraged), as participants would utilize higher response thresholds in the former.

#### 6. Slow errors

Previous studies have often shown that average decision times are longer for inaccurate versus accurate responses. As explained in point 3 above, the cDDM yields this pattern of results, which we also confirmed via simulation.

#### 7. Fast errors

Certain previous studies demonstrated that average decision times are*shorter* for inaccurate versus accurate responses, when the choice difficulty is low and particularly when speedy responses are encouraged. Using simulation analysis, we found that the cDDM can recreate this pattern. However, we note that recreating this pattern while simulating the cDDM required setting the diffusion noise parameter higher than that used to demonstrate the other results above.

#### 8. Increased resolution in confidence with time pressure

Under conditions of time pressure, it seems that there is an increase in the resolution of confidence (see point 3 above). In essence, this means that there will be a larger gap between the average confidence reported for accurate versus inaccurate responses when participants are required or encouraged to decide more quickly. The cDDM can recreate this effect by increasing the rate at which the confidence threshold collapses when decision-makers are in situations where they would feel pressure to respond more quickly. We verified this via simulation.

## DISCUSSION

It is widely assumed that decision-making consists of an accumulation-to-bound process, during which decision evidence (that one option is better than the other) is accumulated over time until a threshold is met (Gold & Shadlen, 2007). This hypothesis is implicit in many mainstream models of decision-making and in particular the DDM, which has been highly successful in explaining choice and reaction time data across a very large number of studies (Ratcliff et al., 2016), but which requires additional assumptions to be able to simultaneously explain choice confidence. An alternative suggestion is that the brain uses a multiplicative urgency signal^4^ that encourages an end to deliberation and a final choice response without excessive further delay (Churchland et al., 2008; Thura & Cisek, 2014; van Maanen et al., 2016). Computational models have been proposed in which the evidence accumulator is amplified at a rate that increases across time, representing the urgency signal (Cisek et al., 2009; Ditterich, 2006). In the DDM, an amplified drift rate is mathematically equivalent to a diminished threshold (if the noise term is adjusted accordingly), so the purported urgency signal might be functionally identical to the collapsing bound in the optimal DDM. In any case, neither of these models considers online (intra-decision) confidence monitoring to be instrumental to triggering a choice. Here, we advance the alternative hypothesis that what is monitored during a decision is the momentary level of confidence across time, and that the decision threshold thus corresponds to a target level of choice confidence. This proposal is related to that of Lee and Daunizeau (Lee & Daunizeau, 2021), who argued that deliberation time optimizes a form of effort-confidence tradeoff, whereby decision-makers will choose to invest time when they expect to achieve a confidence gain that exceeds the anticipated cost of deliberation effort. Here, confidence is used primarily in a *prospective* manner, in the sense that the decision control system sets the target confidence level based on early (pre-deliberation) value representations, and ignores incoming information obtained during deliberation. In contrast, the cDDM model is *reactive*: incremental mental effort accumulates evidence that drives (partially stochastic) changes in confidence until it eventually reaches a pre-defined threshold, which terminates the process and triggers a choice. Both perspectives share the notion that by investing deliberation time, one should expect to achieve higher confidence than would be achieved by choosing prematurely (see also (Chaiken et al., 1989)). A related idea has been proposed in the realm of risky choice (Navarro-Martinez et al., 2018). Here, evidence samples based on probabilistic utility functions serve as input into a t-test calculation (mean / standard error of the mean) and the decision-maker stops deliberating when the t-score reaches a target confidence level. This work was preceded by theoretical studies of perceptual choice demonstrating that optimal decision thresholds should depend on confidence (Drugowitsch et al., 2012). Importantly, the latter model is both prospective (decision thresholds rely on anticipated confidence gains) and reactive (confidence is updated across deliberation time with respect to current levels of accumulated evidence). However, the validity of these decision control models relies on two potentially limiting assumptions: (i) the mechanism by which accumulated evidence changes value representations is known by the system that controls decisions; and (ii) only two options need to be compared. With the cDDM, we relax these assumptions and suggest that there exists a central confidence-monitoring system in the brain that performs a similar function across a wide variety of decisions. This system effectively decides when to decide and is agnostic about the details of the actual decision about to be implemented. Importantly, this unique confidence system would not need to consider specific knowledge about upstream evidence accumulation processes or downstream decision steps (i.e., the choice declaration or implementation), allowing it to perform its duty irrespective of the current choice context. Previous work has suggested the medial prefrontal cortex (mPFC) as such a confidence-monitoring brain region, as fMRI activation represented confidence scores across a variety of value-based tasks (Clairis & Pessiglione, 2022). Another study found that response confidence was tracked by the ventromedial prefrontal cortex (vmPFC) and a prefronto-parietal network, similarly for both memory-based and perceptual decisions (Rouault et al., 2021). Yet another study found that neural firing rates in the rat orbitofrontal cortex (OFC) tracked confidence during an olfactory discrimination task (Kepecs et al., 2008). Under the cDDM, the cessation of deliberation would be influenced not only by confidence accumulation, but also by the rate at which the response threshold collapses. A separate brain region might thus monitor the investment of cognitive effort, with the dorsomedial prefrontal cortex (dmPFC) having been suggested as a potential candidate region (Clairis & Pessiglione, 2022). The interplay between these regions would thus control the dynamics of an effort-confidence tradeoff in deciding when to decide (Lee & Daunizeau, 2021).

We argue that the novel “confidence accumulation” proposal has computational advantages over the standard “evidence accumulation” point of view, and that it permits us to recast several existing empirical findings in novel terms. Furthermore, our approach allows us to make some novel empirical predictions, which we discuss below.

### Computational advantages of the cDDM

From a computational perspective, one may ask, “What is more worthwhile to accumulate in diffusion decision models: evidence or confidence”? Accumulating confidence rather than evidence entails three main computational advantages.

First, the cDDM includes a built-in mechanism that permits accounting for not only the evolving balance of evidence about the values of the different options, but also a measure of certainty about those values. This allows for a more direct readout of confidence at the time a choice is made, relative to the basic DDM (which ignores the concept of value certainty), by taking into account the *precision* of the posterior value estimates. Moreover, it also allows for a direct readout of confidence at any time prior to the choice response (i.e., during deliberation). Thus, the decision apparatus can distinguish from situations where the instantaneous balance of evidence is equal but the precision of that evidence differs, which would allow for the system to optimally determine whether it should continue to deliberate or not. Specifically, in situations where the accumulating evidence has lower precision, deliberation should persist for longer (cf. (Lee & Usher, 2021)). This feature is shared with the model of Drugowitsch and colleagues (Drugowitsch et al., 2012), which is also controlled by a moment-by-moment assessments of confidence.

Second, a decision mechanism that accumulates confidence (like the cDDM) generalizes more easily to decisions between any number of options, compared to one that accumulates evidence (like the basic DDM). The DDM was originally conceived to model decisions between two alternatives. While it could be extended to account for decisions between multiple alternatives, such an extension would require introducing multiple decision variables, which might become implausible as the computational cost would increase exponentially with the number of choice options (Churchland et al., 2008; Roxin, 2019). Additionally, this would be computationally expensive, as it would require keeping track of the momentary rankings across options. On the contrary, the cDDM can model multi-alternative choices using a single compound decision variable that tracks the difference between the posterior probabilities of the best option and all of the other options (see Appendix A). As such, the cDDM can handle arbitrarily-large choice sets without adding much complexity to the computations involved. This is crucial to the extent that real-life decisions typically involve more than two choice options (in contrast to contrived laboratory decisions).

Finally, specific settings of the cDDM will yield decisions that optimally trade off confidence gain (towards the target level) and effort (to increase confidence). This has many implications. First, consider situations where an excessive amount of time is required to increase confidence or when time is lapsing without any increase in confidence: it may then be better to decide immediately (even randomly) rather than to spend additional time deliberating. To afford this optimal cost-benefit tradeoff, the brain needs to be able to prospectively anticipate the confidence gain it could achieve by investing additional deliberation time. In the DDM framework, it can be shown that achieving such optimal decision times eventually implies setting a threshold that collapses with a rate that increases with the cost of deliberation time (Fudenberg et al., 2018; Tajima et al., 2016). This directly applies to the cDDM. Importantly, however, the establishment of this optimal decaying threshold relies upon restrictive assumptions regarding how evidence is used to modify uncertain value representations, which enable the prospective evaluation of the costs and benefits of waiting versus deciding now (Drugowitsch et al., 2012). In particular, these assumptions include the notion that evidence is itself assimilated in an optimal (Bayesian) manner, which neglects systematic errors such as confirmation and/or optimism biases (Kappes et al., 2020; Rollwage et al., 2020; Sharot, 2011) or asymmetries in the impact of evidence for default versus alternative options (Feltgen & Daunizeau, 2021; Lopez-Persem et al., 2016). Under this view, optimal decision timing requires specific decision control systems that cannot generalize to different types of decisions. Nevertheless, one can relax these assumptions by simply considering that the potential magnitude of the cumulative (random) perturbations of value representations will increase with decision time (Lee & Daunizeau, 2021). This would endow the cDDM with a prospective threshold-setting mechanism without committing to detailed assumptions regarding evidence assimilation. In turn, the same decision control system (extended cDDM) would strike at a cost-benefit balance that generalizes over decision types.

Second, consider situations in which decision-relevant information processing can be accelerated, for example, by investing more attentional and/or mnesic resources. This implies that both speed and accuracy can be increased, at the cost of *intensifying* mental effort. This can be achieved if the brain monitors another variable throughout the deliberation process: a resource expenditure monitor^5^ that represents the total amount of cognitive resources that have been invested in the current decision across time. As resource expenditure approaches some critical level, where the anticipated benefit of further expenditure diminishes towards the anticipated cost, this signal would encourage the decision process to terminate even if the target level of confidence were not yet achieved ((Zenon et al., 2019); see (Shenhav et al., 2017) for a review). The ensuing effort-confidence tradeoff would thus modulate optimal decision times with respect to the neurocognitive demands of the decision task. More precisely, optimal decision times would now depend upon the relative costs of effort duration and intensity.

### Reinterpreting Empirical Findings through the Lens of the cDDM

Whether confidence is monitored during (or in parallel with) a choice or after it has been the object of a longstanding debate (Fleming & Daw, 2017). However, many recent studies suggest that confidence judgments influence the development of choice responses rather than just being post-hoc readouts (Schulz et al., 2021). For example, people who already feel confident about their choice after accumulating a certain amount of evidence do not accumulate additional evidence (or they process only choice-congruent information from that point forward; (Rollwage et al., 2020)). Furthermore, different people have different rates of urgency during a choice (which can equivalently be interpreted as having different rates of threshold collapse, meaning that they are more or less willing to accept lower confidence in exchange for saving effort; (Hauser et al., 2017)). All these studies (and others) suggest that the monitoring of confidence is an integral part of the decision process and influences it in real time. One possible way to conceptualize these findings is that the decision process consists of two parallel processes: an evidence accumulation process that is responsible for the choice and a confidence monitoring process (carried out with a slight delay, since it needs to take as input the momentary state of evidence accumulation) that can influence the evidence accumulation (e.g., by setting collapsing bounds). But another possibility is encapsulated by our cDDM model, which simply proposes that decision-making control only requires a single (confidence accumulation) process.

Beyond real-time confidence monitoring, there is already some strong evidence that people explicitly use confidence to control the speed-accuracy trade-off for their choices. In a recent study, participants were asked to make their choices (distinguish between Gabor patches of different orientations) such that they would expect to achieve 70%, 85%, or 90% accuracy across trials (Balsdon et al., 2020). Participants were indeed able to tune their behavior in this way, decreasing RT for lower target confidence and increasing RT for higher target confidence, with subjective confidence levels aligning with objective accuracy percentages. Furthermore, when allowed to respond without any externally-imposed confidence target, participants generally stopped accumulating evidence much earlier and achieved accuracy/confidence levels that were lower than what they were demonstrably capable of (in the high target confidence condition). At the very least, this supports the claim that decision-makers trade confidence against the cost of deliberation time, which is consistent with the cDDM framework. Additional empirical evidence in support of the notion that decision-makers typically continue accumulating evidence until a target level of confidence has been reached was previously shown in the realm of probabilistic decision-making, where participants paid a cost to sequentially reveal cues but stopped whenever the probability of knowing the correct answer rose above a (participant-specific) threshold (Hausmann-Thürig & Läge, 2008). Notably, the same behavior held even in a condition where participants knew that additional cues could not change the relative rankings of the response options, suggesting that increasing confidence was itself an independent goal.

At the neurobiological level, neural signals reflecting (or necessary for) confidence accumulation have been identified during perceptual decisions in parietal (LIP) areas (Kiani & Shadlen, 2009). The fact that this representation is found during the decision period and not (only) afterwards suggests that confidence is not a post-hoc computation but can be integral to the choice. The task used by these authors consisted of a standard “left or right” decision, but to test the effects of confidence on the choice, it also included a “post-decision wagering” condition in which monkeys could select a lower-reward but “sure” offer. The results show that monkeys indeed choose the sure offer when they are (or should be) less confident about the main (“left or right”) choice. While Kiani and Shadlen appeal to a standard evidence-accumulation model to explain LIP firings, what they actually use to fit the behavioral results is an “extension” of evidence accumulation: a model where the standard decision variable is replaced by the log-odds of a correct decision, which (similar to the cDDM) is a representation of choice confidence (Kiani & Shadlen, 2009). This leaves us with the slightly uncomfortable situation where the LIP area is responsible for the “left” versus “right” choice (which is empowered by accumulating evidence) but some downstream process is necessary to calculate the log-odds to make the “sure” choice. A more parsimonious explanation would describe the entire choice (between the three options) as a confidence accumulation process, rather than as an accumulation of evidence followed by a downstream computation of confidence.

Along similar lines, it is possible to reformulate several previous findings that were initially interpreted as evidence for the DDM in terms of the cDDM. For example, a study of multi-alternative choice showed that LIP neuron firing rates gradually increase over deliberation time in a way the resembles evidence accumulation in the DDM (Churchland et al., 2008), which could be difficult to distinguish from confidence accumulation in the cDDM. However, that same study revealed that the average LIP neuron firing rate after target presentation but before motion onset is lower for choices between four alternatives compared to those between two alternatives (Churchland et al., 2008). This is consistent with a neural signal for confidence, since the *a priori* probability of the correct response is lower (25% versus 50%). If the signal was instead monitoring raw evidence, it should be initialized at the same baseline level (representing zero evidence before the start of the trial) regardless of how many options were available to choose from.

### Clearing the empirical hurdles

Pleskac and Busemeyer (Pleskac & Busemeyer, 2010) outlined a number of previous empirical findings that they classified as “hurdles” that any complete model of choice confidence should clear. The cDDM clears seven of the hurdles with no issue. However, we would like to highlight two specific points.

First, Hurdle 7, where inaccurate responses on easy, time-pressured trials generally have lower decision time compared to accurate responses, was not trivial for the cDDM to replicate (e.g., the diffusion noise parameter had to be increased beyond the range used to demonstrate the primary model predictions). We do not consider this to be a major problem, for several reasons. First, this effect has only rarely been reported in prior studies, so its robustness has not been proven. Second, the effect applies to only a very limited percentage of trials, for it specifically addresses inaccurate responses on easy trials. Third, beyond the fact that this effect is limited to easy trials, it is said to be most prominent in conditions of time pressure, further reducing its general applicability. We note that Ratcliff and Rouder (Ratcliff & Rouder, 1998) demonstrated that the effect (Hurdle 7) could be replicated by a DDM that includes trial-by-trial variability in the starting point of the evidence accumulator. While mathematically clever, it is not immediately clear why such variability should be expected at the cognitive level. Previous authors have speculated that starting point variability could capture potential residual effects of the evidence accumulation process on preceding trials (Pleskac & Busemeyer, 2010). Accordingly, if we were to add a starting point variability parameter on top of the cDDM as described in this work, the cDDM would also clear Hurdle 7. Although this seems reasonable, more evidence will be needed before it becomes clear that this is truly something that all confidence models should take into account.

Second, it is worth pointing out that the cDDM inherently replicates the effect summarized by Hurdle 6 (where inaccurate responses on difficult trials generally have higher decision time compared to accurate responses), whereas other models rely on the inclusion of an additional parameter that introduces variability into the drift rate (Pleskac & Busemeyer, 2010; Ratcliff & Rouder, 1998). It is suggested that this variability could capture potential lapses in attention or memory retrieval (Pleskac & Busemeyer, 2010). Again, this seems like a reasonable assumption, though perhaps not a mandatory one.

### Future work to adjudicate between different models

In this theoretical note, we propose an alternate interpretation of traditional evidence accumulation modes of choice. However, since the mathematical formulations of both the cDDM and standard DDMs are fundamentally similar, it is not entirely straightforward to empirically demonstrate the superiority of one over the other. Nevertheless, we believe that such an adjudication could be achieved through novel experiments that test the idea that what is directly monitored during choice deliberation – and what determines when to respond – is confidence (as opposed to “evidence”). One possible way that this could be done would be to train a decoder to transform EEG or MEG activity, measured at the time that participants report their confidence ratings, to readout confidence (across trials). One could then apply the decoding weights at each time point (within trials) and read out the theoretical time-varying (within-trial) confidence dynamics. If this technique works well, one would expect to find continuous, increasing (perhaps fluctuating) trajectories across time that might resemble our illustrative examples in Figure 5. One could perform a similar analysis based on some measure of raw evidence rather than confidence and compare the decoding success, as well as the form of the anticipated trajectories, across confidence and evidence. In principle, this method should be able to provide strong evidence for or against the cDDM relative to the standard DDM.

## CONCLUSION

In this work, we argue that the deliberation process for simple types of decisions (such as two-alternative forced choices based on subjective value) is driven by a confidence monitor rather than a monitor of accumulated evidence in favor of one option over the other. This parsimonious account describes the choice and confidence process as being tightly linked, and it can account for a wide array of behavioral and neural phenomena from the literature. It is also conceptually appropriate for real-world decisions (rather than externally-assigned laboratory tasks). In real life, people will need to set their own personal aspiration levels for choices they make (Simon, 1957). In other words, when faced with multiple options, people not only need to decide but they also need to decide *when* to decide. The confidence-driven drift-diffusion model (cDDM) that we present in this work offers a solution for how people sense the evolution of their confidence across deliberation time and make their choices in a way that optimizes the tradeoff between confidence and mental effort (Lee & Daunizeau, 2021).

## Open Practices Statement

This is a theoretical study. As such, there are no publicly available data or materials, and there were no preregistered experiments.

## Code Availability

The primary analysis code for this study is available at the Open Science Forum at https://osf.io/4yedz/.

## Appendix A

Formulation of confidence for multi-option choices

Define choice confidence *P*_*c*_ as the probability that the (predicted) experienced value of the (to-be) chosen option is higher than that of all of the (to-be) unchosen options. Critically:

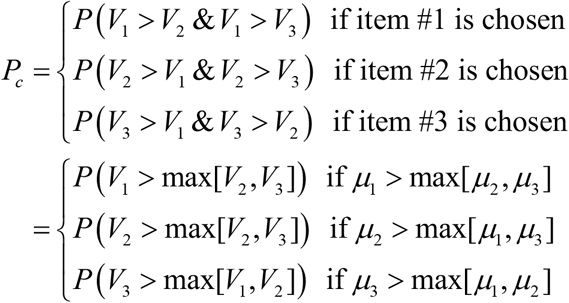

where the second set of equations derives from assuming that the option with the highest expected value will be chosen. Choice confidence is thus derived from a comparison between the best option (i.e., the option with the highest expected value μ_*best*_) and the maximum of the rest of the options. For example, for a choice set containing three options, with the first option having the highest expected value, the expected maximum value of the other options is (Nadarajah & Kotz, 2008):

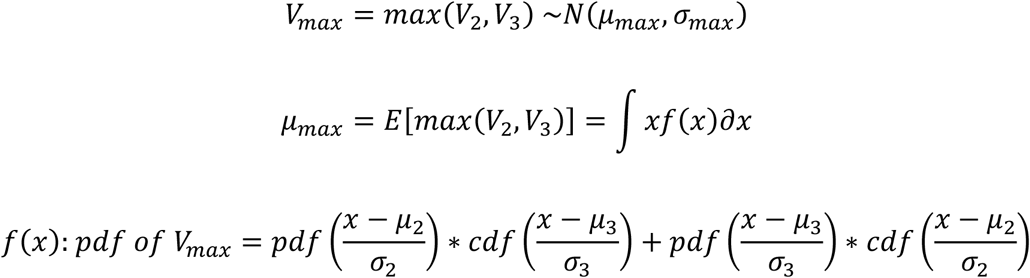

For a choice set with four options, with the first option having the highest expected value, an iteration of this procedure will provide the expected value of the maximum of the options other than the best:

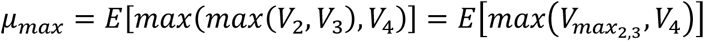

The first and second moments of the maximum function can be calculated iteratively to obtain first the expected maximum value of two of the three non-best options, then the expected maximum value of this value and the third non-best option:

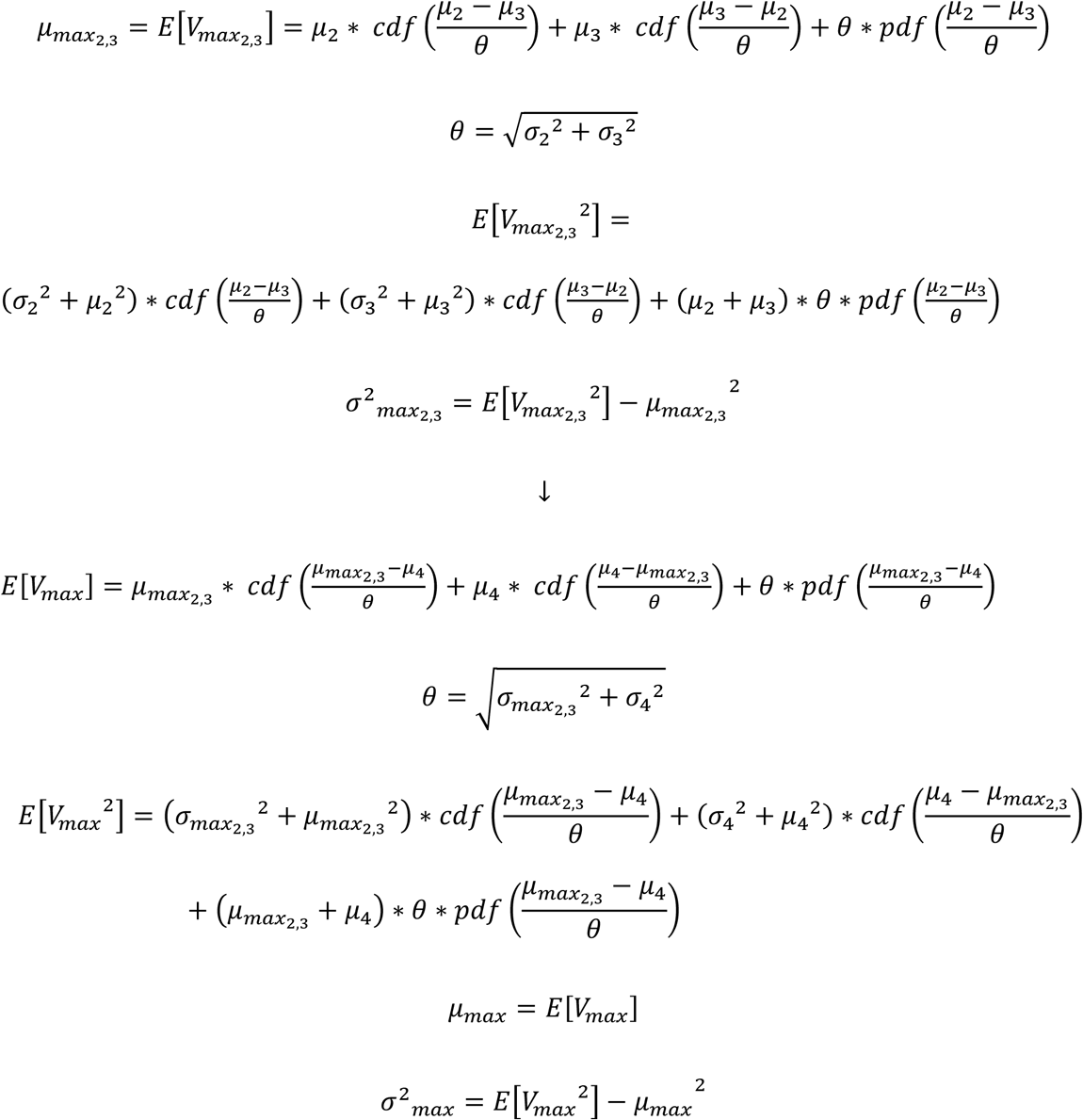

Clearly, this iterative process could be repeated indefinitely, allowing for a choice option set of arbitrarily large size.

Once the expected maximum value of the non-best options is determined, the DM will compare this value with the expected value of the best option. Thus, the decision variable (DV) will be a normally distributed random variable with mean equal to the difference of the best and max-non-best means and variance equal to the sum of the best and max-non-best variances:

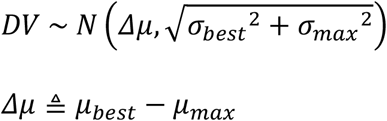

Subjective uncertainty regarding option values eventually translates into uncertainty regarding what the “correct choice” is. This is summarized in terms of the probability P_c_ of “committing to the correct choice”:

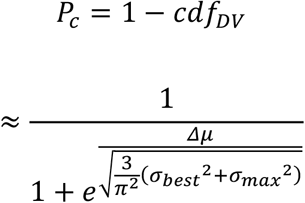

where *cdf* is the cumulative density function of the corresponding normal distribution, evaluated at *DV* = 0, and the second line derives from a simple sigmoidal approximation to the normal cumulative density function (Daunizeau, 2017b).

## Appendix B

Non-decision time and starting point bias

In a possible extended version of the model, the cDDM assumes that during the pre-deliberation NDT, when the options are being perceived and identified, a low-precision value representation of the options is automatically formed (Lee & Daunizeau, 2021). Previous studies have shown that the brain automatically starts to process information related to value even when not actively deliberating about it (i.e., outside of actual decision time) or when preparing to deliberate about something unrelated (Lebreton et al., 2009, 2015; Lopez-Persem et al., 2020). For the purposes of our model and theory, NDT corresponds to the time required for the perception and identification of the options (we choose not to consider post-deliberation motor response NDT). Crucially, the cDDM assumes that during this period, value representations of the options are formed in parallel to perceptual processes, but with rather low precision. This is because in this early sensory processing stage, the options are not yet completely perceived or identified. This early representation formation can be formalized in the same manner as during explicit deliberation, only with lower precision (σ_NDT_^2^ > σ^2^; e.g., lower attention will be assigned to the valuation task at this stage):

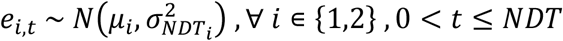

At the end of NDT, the decision-maker will have an initial estimate of the value (v) of each option, as well as an initial estimate of precision (p) for the value estimates:

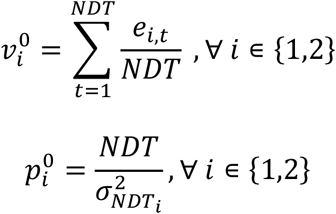

Based on the initial estimate and precision of value for each option, the decision-maker will already have an initial sense of confidence by the time deliberation commences. Thus, whereas the confidence monitor (c) begins the decision process at zero, it typically arrives at a non-zero point by the end of the NDT. This effectively creates a *starting point bias* (z) equal to the subjective probability that one of the options is better than the other (i.e., confidence) prior to explicit deliberation:

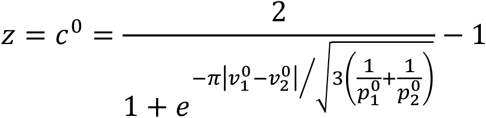

More generally, both the momentary value estimates and the momentary confidence level will continue to evolve across deliberation time. These variables will incorporate both the information that was automatically processed during NDT and the information that was deliberately processed after NDT:

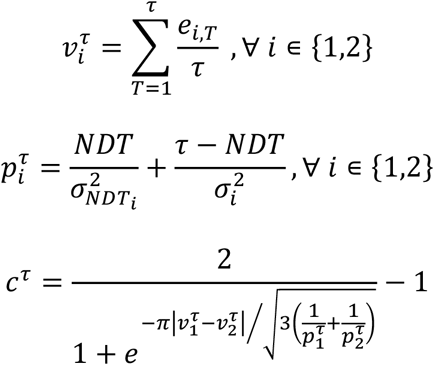

where τ is the time index at the moment the choice response is entered.

## Footnotes

1 This is particularly true in the domains of perceptual and preferential decision-making, although it is also becoming more common in the domain of economic decision-making.

2 The collapsing threshold is a hallmark of the optimal rendition of the DDM, though it is not a feature of the standard DDM. However, some non-optimal versions of the DDM have also included collapsing thresholds (Voskuilen et al., 2016).

3 For our purposes here, we will not consider non-essential parameters such as x_0_ and NDT. For a possible extension of our model that might relate to x_0_ and NDT, see Appendix B.

4 Urgency signals are typically presented with respect to time specifically, where responses are made sooner in order to save time. Rising deliberation costs and urgency are not mutually exclusive, and in fact the latter can be considered a special case of the former. Furthermore, one could monitor both time urgency and other costs of deliberation, and diminish the confidence threshold in response to either or both.

5 There is debate about what resources are consumed in a costly manner during cognitive processes, including metabolites, time, or capacity (Zenon et al., 2019; see Shenhav et al., 2017 for a review). For our purposes, we remain ambivalent and use the term resources in a more abstract sense (i.e., whatever the true costly resource turns out to be, our model will not change).

